# Viral soluble decoy receptor as a novel treatment for IFN-I triggered diseases

**DOI:** 10.1101/2025.08.12.669887

**Authors:** Francisco Javier Alvarez-de Miranda, Isabel Alonso-Sánchez, Pilar Delgado, Antonio Santos-Peral, Bruno Hernaez, Antonio Alcamí

## Abstract

Besides its antiviral action, type I interferon (IFN-I) plays a key role in triggering the inflammatory response and immunopathology in a group of autoinflammatory disorders known as interferonopathies. Current therapies for these diseases focus on blocking IFN-I signaling at different levels, but with incomplete success. Throughout evolution, to counteract the host immune response, poxviruses have developed decoy receptors, soluble proteins secreted from infected cells during infection that bind and block key host cytokines. One of these, the poxvirus IFN-I binding protein (IFNαβBP) is a unique soluble receptor with the ability to block many IFN-I subtypes with broad species specificity. In addition, this protein interacts with glycosaminoglycans on the cell surface while binding IFN-I, which enhances its immunomodulatory potential by allowing the retention of this receptor around the site of infection. In this study, we have deepened into how the poxvirus IFNαβBP modulates IFN-I to apply this knowledge to the development of new anti-IFN-I therapeutics. With this purpose, a set of IFNαβBP-based recombinant proteins were generated and tested for their ability to neutralize IFN-I and bind to the cell surface *in vitro*, and their immunomodulatory action was tested in the mousepox model of pathogenesis. Finally, the therapeutic potential of some of these selected viral IFN-I inhibitors was validated in two different murine models of autoinflammatory disease, imiquimod-induced psoriasis and pristane-induced lupus.

## Introduction

Type I interferons (IFN-Is) are a family of cytokines with pleiotropic functions on innate and adaptive immunity, including the induction of cell programs that increase resistance to viral infections. Their biological activity starts with the binding to heterodimeric cellular receptors consisting of a high affinity receptor chain IFNAR2 and a low affinity receptor chain IFNAR1, with affinities in the nM to μM range, respectively (1–3). Whereas the extracellular portions of the IFN-I receptors bind their cognate ligands, the intracellular domains constitutively associate with Janus Kinases (JAKs) ) (4,5) that initiate IFN-mediated intracellular signaling cascades through phosphorylation and recruitment of signal transducers and activators of transcription (STATs). This ultimately leads to the activation of IFN-stimulated gene factor 3 (ISGF3) and the expression of IFN stimulated genes (ISGs) (6) that target conserved aspects of viral infection (7,8). The various immune functions of IFN-Is have made them very promising agents for the treatment of several diseases like cancer and viral infections (9).

However, IFN-I benefits do not come without a cost. Treatment with exogenous IFN-I leads to various adverse effects for a few hours after administration (10), illustrating how IFN-I dysregulation can negatively impact health. In addition, increased levels of endogenous IFN-I can play a key role in triggering the inflammatory response and the pathology in disorders known as interferonopathies (11), a set of mendelian inborn errors of immunity that are comprised in the broader group of autoinflammatory disorders (12). At least 38 Mendelian genotypes with mutations in nucleic acids sensing, IFNAR signaling, mitochondrial integrity or proteasome can lead to type-I interferonopathies in which IFN-I is directly or partly responsible for the pathology (11). These usually present during infancy and lead to severe clinical manifestations or even death, like the case of Aicardi-Goutières Syndrome (AGS), the prototypical type-I interferonopathy (13).

Treatment approaches for AGS have focused on blocking IFN-I at different levels, like enhancing the removal of self-nucleic acid stimuli with reverse transcriptase inhibitors (14), limiting IFN-I production by inhibiting JAK1 (15) or blocking IFN-I signaling with anti IFNα antibodies (16). Regrettably, there are no officially approved drugs for the treatment of AGS yet (17), leading to a constant need for new therapeutic approaches. IFN-I has also attracted attention as one of the key factors in the pathogenesis of systemic lupus erythematosus (SLE). This disease is characterized by an increased amount of apoptotic material (DNA, RNA and nuclear proteins) that is recognized by plasmacytoid dendritic cells (pDCs) and induces the secretion of IFN-I, leading to an immune dysregulation that causes a major life threat for patients (18–21). Traditional therapeutic approaches for SLE include nonspecific anti-inflammatory drugs (22,23) and targeting B cells (24). However, the extensive links between SLE and IFN-I signaling (25,26) have raised interest in new therapeutic approaches targeting IFN-I.

In the treatment of other autoinflammatory diseases, soluble versions of cellular receptors have been used to antagonize exacerbated proinflammatory cytokines that cause pathology, such as the soluble TNFα receptor Etanercept for the treatment of rheumatoid arthritis (27). However, no soluble versions of IFNAR have been developed as therapeutics, probably due to the complexity of this heterodimeric receptor and the diversity of IFN-I ligands.

Viruses are obligate parasites whose survival fully depends on their ability to replicate in hostile environments. Their diversity, fast life cycle and adaptation capacity have made them come up with unique biological mechanisms during their millions of years of co-evolution with their hosts. In particular, Poxviruses are a family of large DNA viruses that have developed a wide array of strategies to counteract the antiviral action of IFN-I during infection, from its induction to its effector functions (28,29). These include a unique viral mechanism: the blockade of soluble IFN-I with a secreted decoy receptor, the viral IFN-I binding protein (IFNαβBP).

The poxvirus IFNαβBP is a soluble decoy receptor, secreted from infected cells to bind IFN-I and prevent its recognition by cellular cognate receptors, thus impairing antiviral signaling (30,31). Interestingly, the calculated affinities of the IFNαβBP for some IFN-Is are around 0.1 nM, even higher than that of IFNAR2 (32). Besides, in contrast to the high species specificity of cellular IFN-I receptors, the poxvirus IFNαβBP binds different IFN-I subtypes with high affinity and IFN-I from different species including human, murine, rabbit and bovine (31–33). It has been suggested that its high inhibitory potency and broad specificity are related to its unique structure, which is curiously not related to the fibronectin-III like domains of the cellular IFNAR but rather composed of three immunoglobulin-like domains. After secretion, the IFNαβBP binds to the cell surface of infected and uninfected cells, where it retains the ability to neutralize IFN-I (34). The interaction with the cellular surface is independent from the ability to block IFN-I and is mediated by a glycosaminoglycan (GAG) binding domain (GAGBD) located in the first immunoglobulin-like domain (D1) (35). This cell surface binding ability has proven to be essential for effective immunomodulation, as shown by the severe attenuation observed after mice infection with vaccinia virus (VACV) and ectromelia virus (ECTV) expressing an IFNαβBP lacking cell surface binding properties (36). It has been suggested that poxviruses could have developed this cell surface binding to retain the secreted decoy receptor near the infected cells, a strategy to increase the local concentration of the protein and improve its immunomodulatory action. Additionally, some studies have hinted that the ability to block IFN-I could be mostly associated with the last immunoglobulin-like domain (D3) (34,37), which only represents a third part of the full protein.

The possibility of learning how pathogens modulate the immune response as a source for new drugs has raised interest in academia and biotech companies (38,39). Virus-encoded strategies to effectively counteract host immunity might constitute an extensive natural repository for new immune-modulating molecules that can help us in the fight against disease conditions caused by an uncontrolled immune response. In particular, the poxvirus IFNαβBP has already been used to neutralize the IFN-I response during RNA mediated gene delivery (40,41) or cell reprogramming into stem cells (42). Additionally, this protein was tested as a potential treatment in pilot studies to alleviate IFN-I-mediated neurotoxicity during HIV encephalitis in mice (43,44). In this study, we explored the IFNαβBP potential as a therapeutic tool and tested it for the first time in two different murine models of autoinflammatory disease, which aligns with the need for new therapeutic approaches to block IFN-I in the treatment of type-I interferonopathies.

## Results

### A minimal domain from the poxvirus IFNαβBP can bind and block various subtypes of IFN-I

The crystalized structure of the complex between murine IFN-α5 and the IFNαβBP from ECTV (Moscow strain, PDB:3OQ3) was used as a template to study the interaction of VACV IFNαβBP with IFN-I. This allowed to generate a computational model showing the three domains (D1, D2, D3) **(Fig.1 A)** and the positively charged region of the GAGBD that interacts with the negatively charged cell surface GAGs **(Fig. 1 B)**. In addition, a stable saline bridge between the IFNαβBP D3 Asp276 and the IFN-α5 Arg150 was observed, and the D3 Asp278 was close to the key IFN-α5Arg33 **(Fig. 1 C)**. During cellular recognition, IFNAR2 makes extensive interactions with the conserved Arg33 (45–47) and the Met148 and Arg149 (48) of the IFN-I protein (IFNα2 numbering) before the formation of the tertiary complex with IFNAR1. Therefore, in the IFNαβBP-IFN model, D3 is blocking IFN-I by covering the critical region for its recognition by the high affinity cellular receptor IFNAR2.

**Figure 1.**
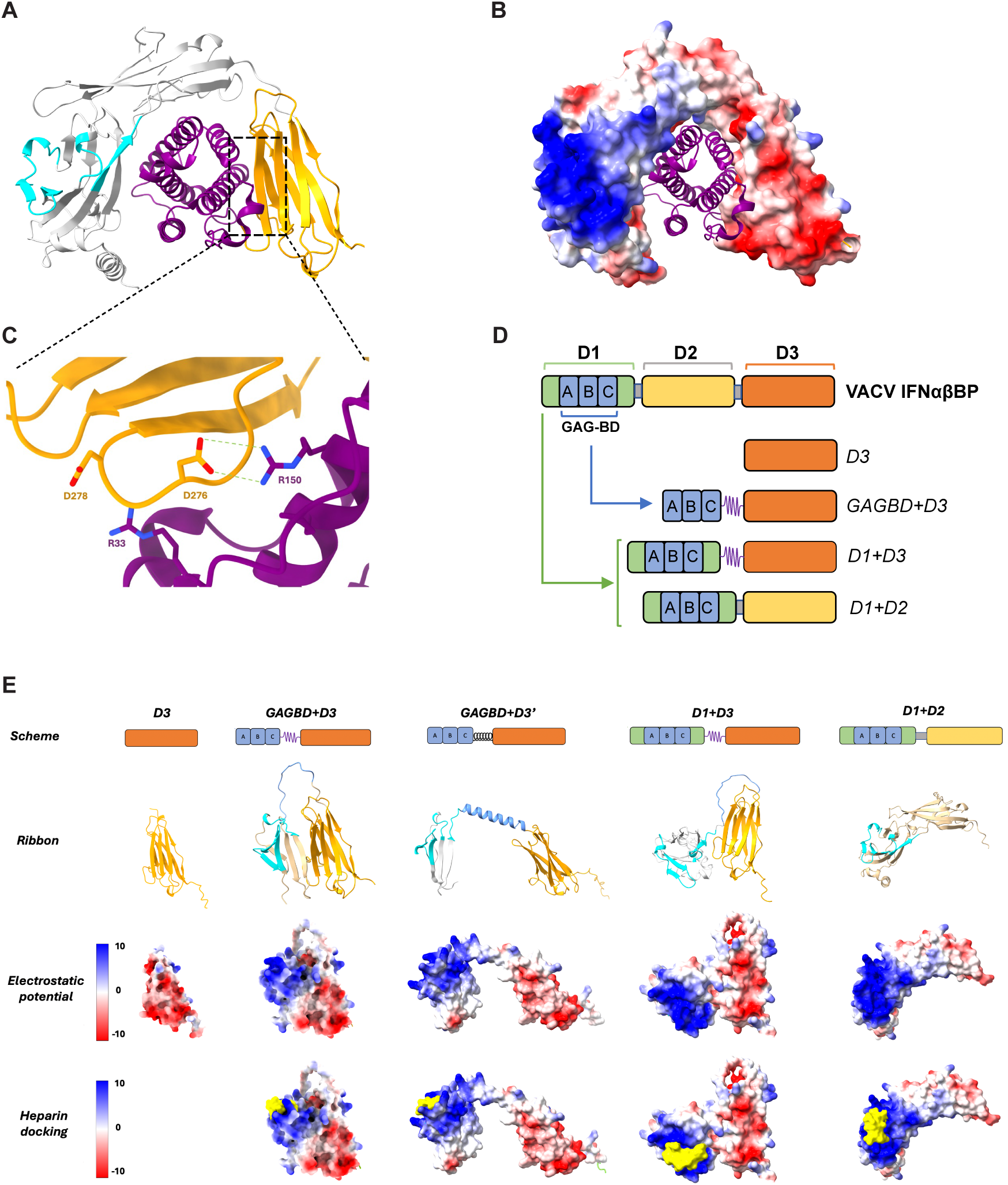
IFNαβBP-based proteins containing D3 bind and block IFN-I. **(A)** Computational model for VACV IFNαβBP, overlapped over PDB:3OQ3 to model interactions with IFN-I, highlighting the GAGBD (cyan) and D3 (orange). **(B)** Coulomb surface potential of the model in Fig.1 A. (**C**) Detailed view of saline bridges between conserved amino acids for the critical interaction between the IFNαβBP and IFN-I. **(D)** Schematic representation of the designed IFNαβBP-based recombinant proteins. **(E)** Alphafold generated models for IFNαβBP-based proteins, with their Coulomb surface potentials and heparin (yellow) docking.

To study the interaction between the IFNαβBP, IFN-I and GAGs, the following set of IFNαβBP-based recombinant proteins were designed and expressed in insect cells (**Fig. 1 D**): 1) the isolated D3, 2) D3 fused to the minimal GAGBD described for the IFNαβBP (35) and 3) D3 fused to the full D1 containing the GAGBD. In addition, a fusion of the first and second domain was expressed as a control construction that should bind to the cell surface without blocking IFN-I. Computational models for these recombinant proteins showed a densely positive surface in the putative cell surface binding domains, and revealed that these surfaces were able to dock heparin molecules, suggesting their potential ability to bind GAGs (**Fig. 1 E)**.

The recombinant proteins were purified (**Fig. S1**) and their ability to bind various subtypes of IFN-I was tested by surface plasmon resonance (SPR) (**Table 1**). All proteins containing D3 showed binding for the IFN-Is assayed, although their calculated binding affinity constants were globally 1 or 2 orders of magnitude higher than that of the native IFNαβBP. Interestingly, despite being the minimal IFN-I binding domain, D3 showed higher affinity for hIFNα compared to the recombinant proteins GAGBD+D3 and D1+D3.

**Table 1.** Affinity constants (nM) for murine and human IFN-Is determined by SPR. NBD: no binding detected.

|  | **mIFNα** | **hIFNβ** | **hIFNα** |
| --- | --- | --- | --- |
| **D3** | **1.96 ± 0.78** | **28.98 ± 4.26** | **2.77 ± 2.38** |
| **GAGBD+D3** | **1.96 ± 0.68** | **61.63 ± 5.45** | **17.41** |
| **D1+D3** | **0.60 ± 0.13** | **47.77 ± 1.99** | **47.5** |
| **D1+D2** | NBD | NBD | NBD |
| **IFN$\alpha\beta$BP** | **0.02 ± 0.002** | **0.09 ± 0.022** | **0.35** |

Although the recombinant proteins were able to bind IFN-I in SPR experiments, the binding of host ligands by viral proteins does not always translate into the blockade of their biologic activities (32,49). To determine their capacity to block IFN-I biological activity, a previously described neutralization assay of the IFN-sensitive vesicular stomatitis virus (VSV) was used (32). Although full-length IFNαβBP was more efficient at low doses, D3, GAGBD+D3 and D1+D3 were able to block mIFN⍺ activity in a dose-dependent manner (**Fig 2 A**). D1+D2 did not show any anti-IFN-I effect, as expected because it lacks the critical D3 domain. Next, the potential blockade of human IFN-Is in human HeLa cells was investigated. The blockade of hIFNβ was similar to that of mIFN⍺ (**Fig 2 B**), with all proteins containing D3 showing a dose-dependent protection. However, when analyzing the blockade of hIFN⍺, the isolated D3 confirmed its SPR-determined superior affinity compared to the other truncated proteins, being the only one that displayed some protective effect at the hIFN⍺ concentrations tested (**Fig 2 C**). As observed for mIFN⍺, full-length IFNαβBP was more efficient at inhibiting the activity of hIFNβ and hIFN⍺ than the truncated proteins.

**Figure 2.**
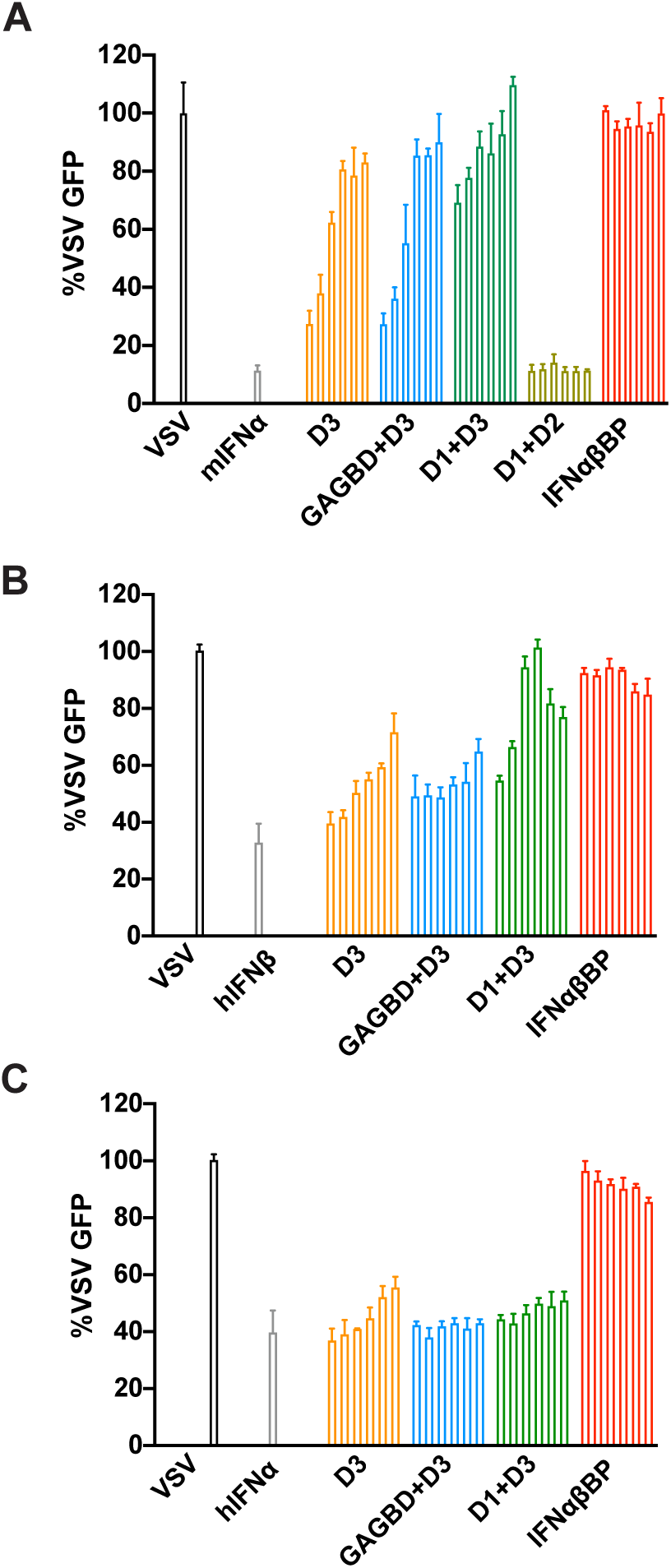
Blockade of IFN-I biological activity by the IFNαβBP-based recombinant proteins. Vesicular stomatitis virus (VSV)-GFP replication, measured as GFP fluorescence, in cells treated with IFN-I. **(A)** VSV replication in L929 cells treated with 0.5 U murine IFN⍺ (mIFN⍺) preincubated with truncated proteins (1, 2.5, 5, 10, 25 and 50 nM) or the full IFNαβBP (0.5, 1, 2.5, 5, 10 and 25 nM). **(B)** VSV replication in HeLa cells treated with 2.5 pM human IFNβ (hIFNβ) preincubated with recombinant proteins (5, 10, 25, 50, 100 and 250 nM). **(C)** VSV replication in HeLa cells treated with 15 pM human IFNα (hIFNα) preincubated with recombinant proteins (5, 10, 25, 50, 100 and 250 nM). A representative experiment of at least three experiments performed is shown.

### The IFNαβBP cell surface binding ability can be transferred to the minimal domain D3

We have previously demonstrated that the IFNαβBP from VACV (35) and ECTV (36) can bind GAGs at the cell surface by interactions with basic regions in the first domain of the protein. Given the importance of the cell surface binding for the IFNαβBP, we proposed to transfer this ability to the minimal IFN-I blocking domain D3 to improve its immunomodulatory activity. We hypothesized that fusion of the GAGBD or the full first domain D1 to D3 could transfer the cell surface binding ability to the minimal IFN-I blocking domain. To test this, equal amounts of D3, GAGBD+D3, D1+D3, D1+D2 and IFNαβBP were incubated with CHO-K1 cells and cell surface binding was quantified by flow cytometry (**Fig. 3 A**). The recombinant receptors displayed a spectrum of affinities for the cell surface, with D3 showing similar binding to a negative control, the unrelated semaphorin VACV orthologue (SEMA). While all the proteins containing the full D1 (D1+D3 and D1+D2) displayed a surface binding pattern similar to that of the full IFNαβBP, GAGBD+D3 showed a drastically reduced binding ability, representing only 10% of the binding of the full protein. A similar experiment was performed to visualize cell surface binding on CHO-K1 cells by immunofluorescence (**Fig. 3 B**). To confirm that interaction with the cell surface is mediated by heparin (35) similar flow cytometry experiments with mutant CHO-745, which completely lack surface GAGs, or CHO-677, with a specific depletion in heparin, were performed. No cell surface binding was observed in neither of the CHO mutant cells, confirming that the interaction is mediated through heparin **(Fig. S2)**.

**Figure 3.**
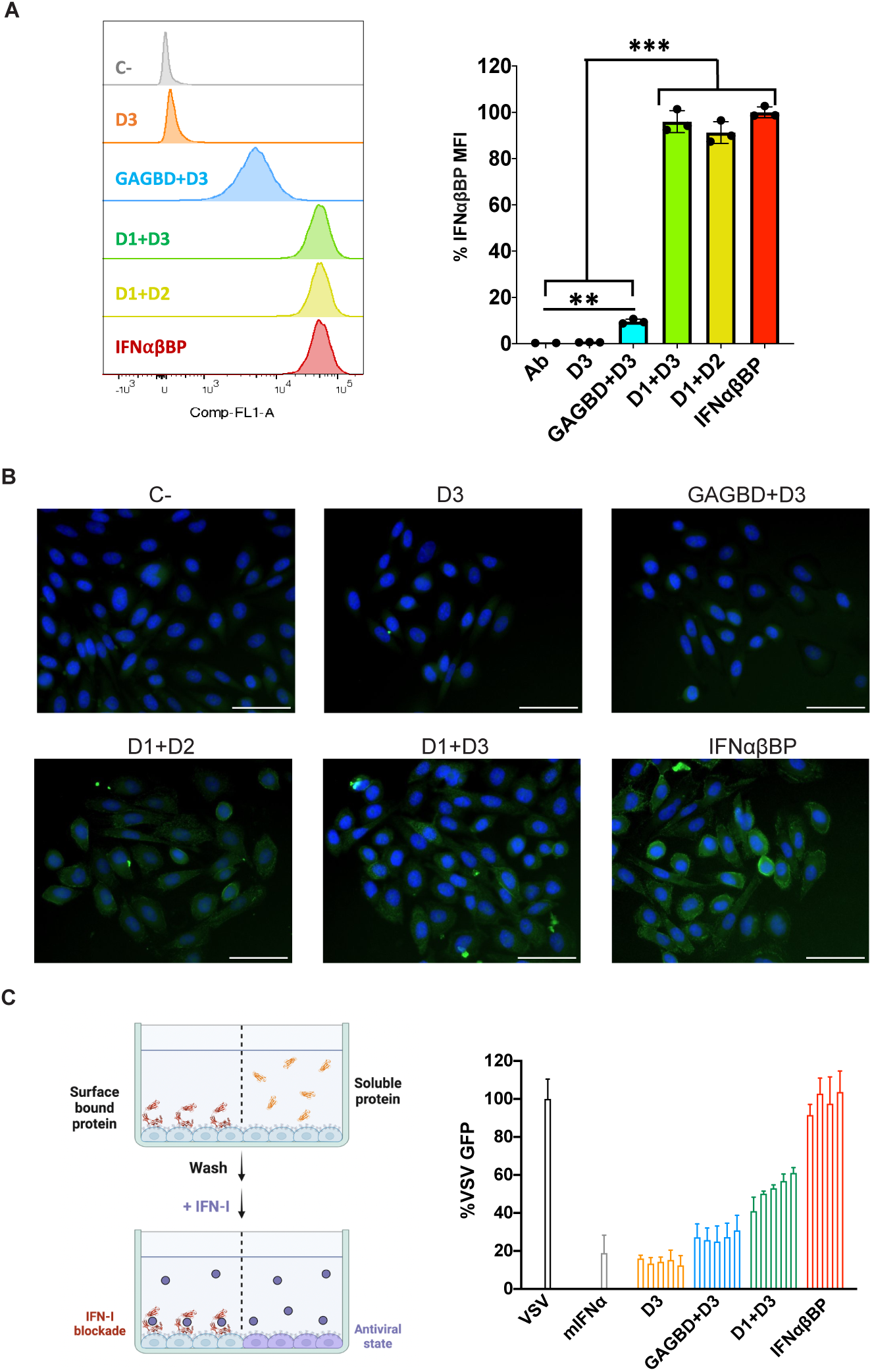
IFNαβBP-based proteins display a spectrum of affinities for cell surface GAGs. **(A)** CHO-K1 cells were incubated with 250 nM of purified recombinant proteins at 4° C, stained for V5 tags and analyzed by flow cytometry. Cell surface binding is shown relative to binding of full-length IFNαβBP. A representative experiment from three experiments performed is shown. **(B)** CHO-K1 cells were incubated with 250 nM of purified recombinant proteins at 4° C and observed under the fluorescence microscope after staining of V5 tags. Scale bar 50 μm. A representative experiment of two experiments performed is shown. **(C)** To determine simultaneous IFN-blocking and cell-surface binding abilities, increasing amounts of truncated proteins (1, 5, 10, 25 and 50 nM) or the full IFNαβBP (1, 5, 10 and 25 nM) were incubated on L929 cells and then thoroughly washed before adding 0.5 U mIFN⍺. Then, cells were infected with 2 pfu/cell VSV-GFP and viral driven fluorescence was measured at 24 hours post-infection (hpi). Results were analyzed with unpaired t test, ** = p<0.01. *** = p<0.001.

Finally, the ability of the proteins to block IFN-I while bound to the cell surface was evaluated in a modified version of the previous assay (Fig. 2), in which unbound proteins are washed away before IFN addition and VSV infection, so only the proteins that remain attached to the cell surface after washing can block IFN-I (**Fig. 3 C**). This experiment was performed with mIFN⍺ and L929 cells, as these conditions gave the best results. As expected, D3 was washed away in this setup. However, although GAGBD+D3 only exerted a small anti-IFN-I effect, D1+D3 confirmed its surface binding capacity and showed dose dependent anti-IFN-I activity despite extensive washing.

### D3 can exert immunomodulatory action during ECTV infection

During ECTV infection of susceptible BALB/c mice, viral immunomodulation leads to a decreased immune response against the virus that translates into greater virulence, making this model suitable to study the action of poxvirus immunomodulators (36,50,51). To study the anti-IFN-I properties of the IFNαβBP-based proteins of interest during mouse infection, we generated recombinant ECTVs in which the native ECTV IFNαβBP (EVN194) was substituted for D3 or D1+D3 (**Fig. 4 A**). After successful insertion of the coding sequences into the virus genome, recombinant ECTVs were fully sequenced at high depth and *in vitro* replication was evaluated. The viral progeny was equivalent to that of parental ECTV WT in both one-step and multistep growth curves, discarding any defects in virus production or spreading *in vitro* (**Fig S3**).

**Figure 4.**
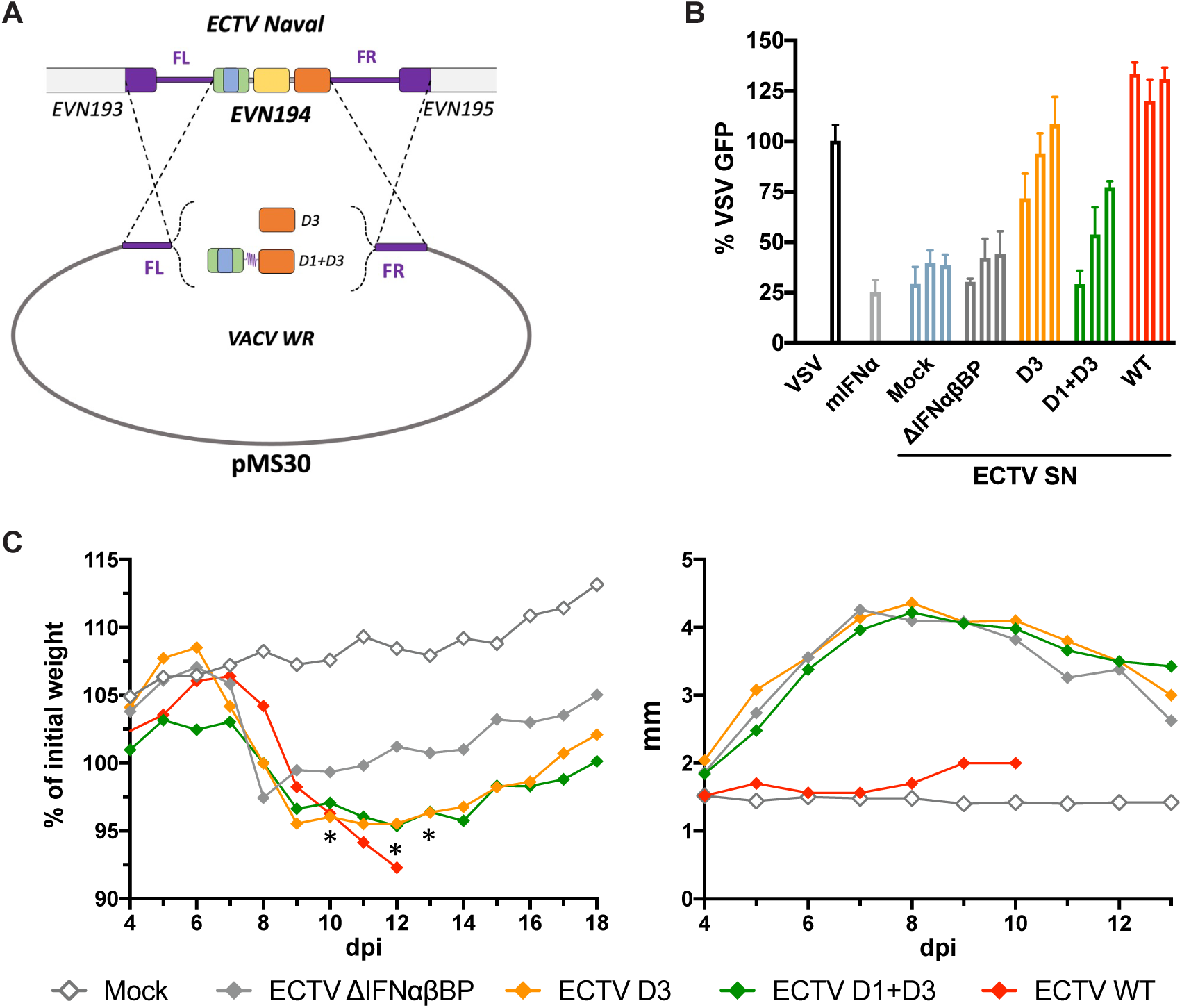
Expression of D3 improves immunomodulation compared to ECTV ΔIFNαβBP. **(A)** Schematic representation of the recombination process for the generation of recombinant ECTVs. **(B)** mIFN⍺ (0.3 U) was preincubated with concentrated viral supernatants (5, 10 or 25 μl) and added to L929 cells before VSV-GFP infection. A representative experiment of two experiments performed is shown. **(C)** Bodyweight evolution and footpad inflammation after subcutaneous inoculation of mice with 10^6^ pfu of recombinant ECTVs or 10 pfu of ECTV WT via footpad (N=5 per condition). Results were analyzed with multiple T-tests corrected with Holm-Sidak method, * = p corrected < 0.05 for ECTV D3 and ECTV D1+D3 compared to ECTV ΔIFNαβBP. A representative experiment of two experiments performed is shown.

To evaluate whether these recombinant viruses expressed functional IFN-I decoy receptors, a new VSV neutralization assay using concentrated supernatants from recombinant-ECTV infected cells was performed. In this setup, ECTV D3 and ECTV D1+D3 supernatants induced better protection against mIFN⍺ than ECTV ΔIFNαβBP or uninfected supernatants (**Fig. 4 B**), proving the expression of active decoy receptors with anti-IFN-I activity in the generated viruses. Interestingly, ECTV D3 supernatants showed better anti-IFN-I activity than those from ECTV D1+D3, hinting at a greater concentration of the protein in this supernatant. This could be explained by D1+D3 being retained on the surface of cells during infection and therefore being less present in the culture supernatant.

Next, the recombinant ECTVs infection in susceptible mice was studied and compared to the infection of ECTV WT and ECTV ΔIFNαβBP (**Fig. 4 C**). Both recombinant ECTVs were highly attenuated compared to ECTV WT but caused higher bodyweight losses than ECTV ΔIFNαβBP. This indicated that D3 and D1+D3 were able to exert some immunomodulation during ECTV infection. However, despite the cell surface binding ability of its encoded decoy receptor, ECTV D1+D3 induced similar virulence to ECTV D3. In addition, no differences were observed in footpad inflammation between ECTV D3, ECTV D1+D3 and ECTV ΔIFNαβBP, indicating the limitations of ECTV infection as a model to study the immunomodulatory potential of these viral IFN-I inhibitors.

### Viral IFN-I inhibitors D3 and IFNαβBP reduced DCs activation in Pristane-induced lupus (PIL) mice

The therapeutic potential of the viral IFN-I inhibitors D3 and IFNαβBP was tested in the PIL mouse model. This murine model is characterized by a lupus-like autoinflammatory disease with a clear IFN-I signature after intraperitoneal injection of pristane (2,6,10,14-tetramethylpentadecane). Although the development of later symptoms in PIL mice can take weeks or even months, IFN-I dysregulation plays a critical role in triggering the pathology from the first days post-injection (52).

PIL mice were treated with IFNαβBP, D3 or PBS for a short period of time in the early days after pristane injection **(Fig. 5 A)**. After the induction of the model and treatment, the immune infiltrate in the peritoneal cavity was studied, focusing on the specific effect of the treatment on DCs. Despite most cell types being IFN-I producers, DCs are known for their ability to rapidly produce large amounts of IFN-I under specific conditions (53), and their key role in PIL has been previously described (54). Although the percentage of DCs (CD45+ CD11c+) remained unchanged under different experimental conditions, a defined population of activated DCs (CD80+ CD86+) appeared after induction with pristane and was significantly reduced upon treatment with D3 or IFNαβBP (**Fig. 5 B**).

**Figure 5.**
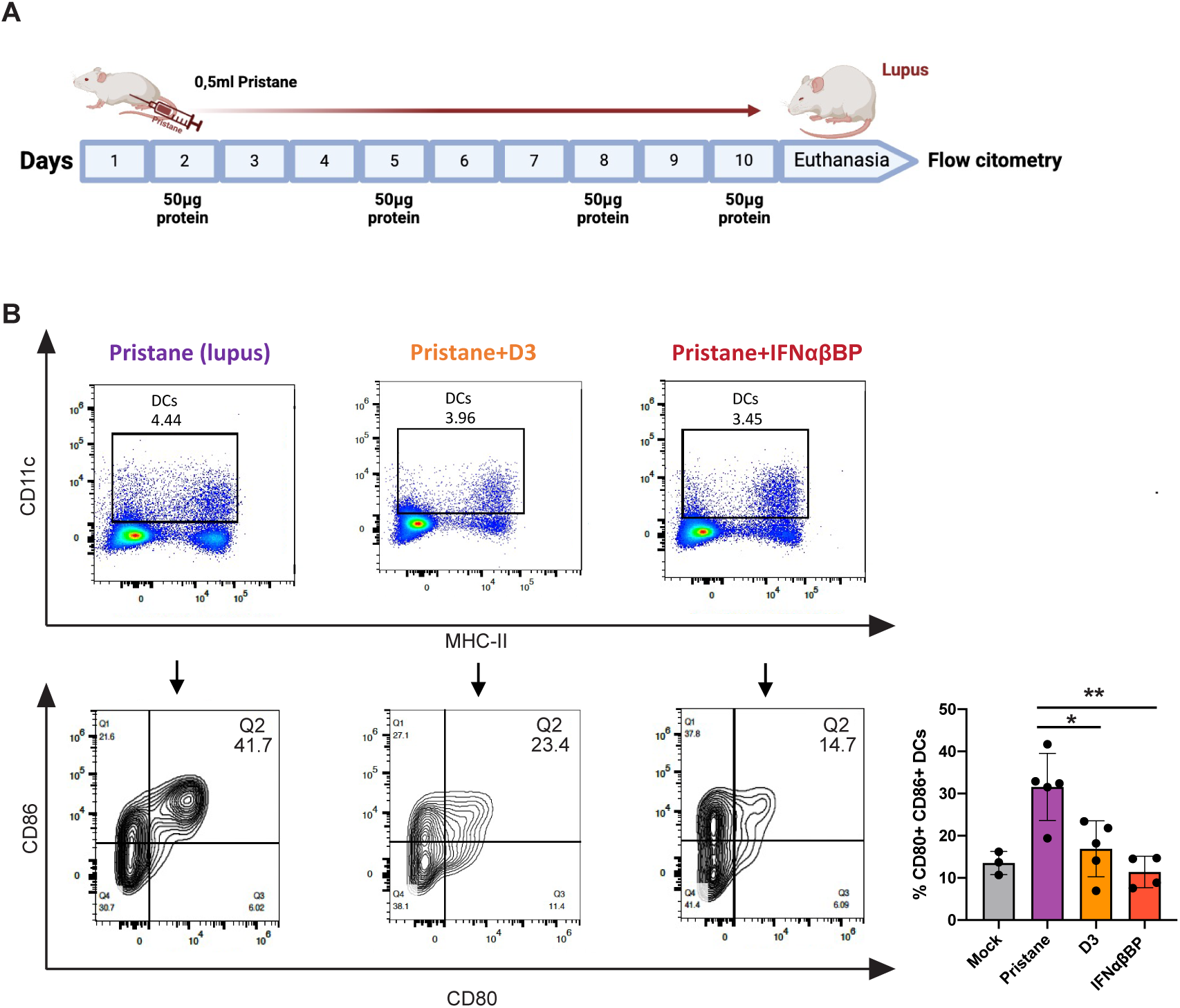
Viral IFN-I inhibitor treatment in a mouse model of PIL. **(A)** BALB/c mice were injected with 0.5 ml pristane and intraperitoneally treated with 50 µg of IFNαβBP, D3 or PBS every 48-72h for 10 days. **(B)** Levels of activated DCs (CD45+, CD11c+, CD86+, CD80+) in the peritoneal cavity. Results were analyzed with Mann Whitney test, * = p<0.05, ** = p<0.01. N was a minimum of 4 mice for the treated groups and 3 for the untreated group. A representative experiment of two experiments performed is shown.

### Viral IFN-I inhibitors D3 and IFNαβBP reduced inflammation and psoriatic epidermal thickening in a mouse model of imiquimod-induced psoriasis (IIP)

The immunomodulatory potential of the recombinant viral IFN-I inhibitors D3 and IFNαβBP was also evaluated in a mouse model of IIP, an extensively used model for autoimmune disease (55). Consecutive topical application of imiquimod in the ears of BALB/c mice results in skin inflammation and psoriasis-like disease (56) through the activation of TLR7 and TLR8 and the induction of IFN-I (57). We hypothesized that viral IFN-I inhibitors could ameliorate pathology in this model. After IIP induction and treatment, one ear was fixed for histological analysis, and the axillar and sub-iliac lymph nodes (LNs) were collected for flow cytometry analysis (**Fig. 6 A**).

**Figure 6.**
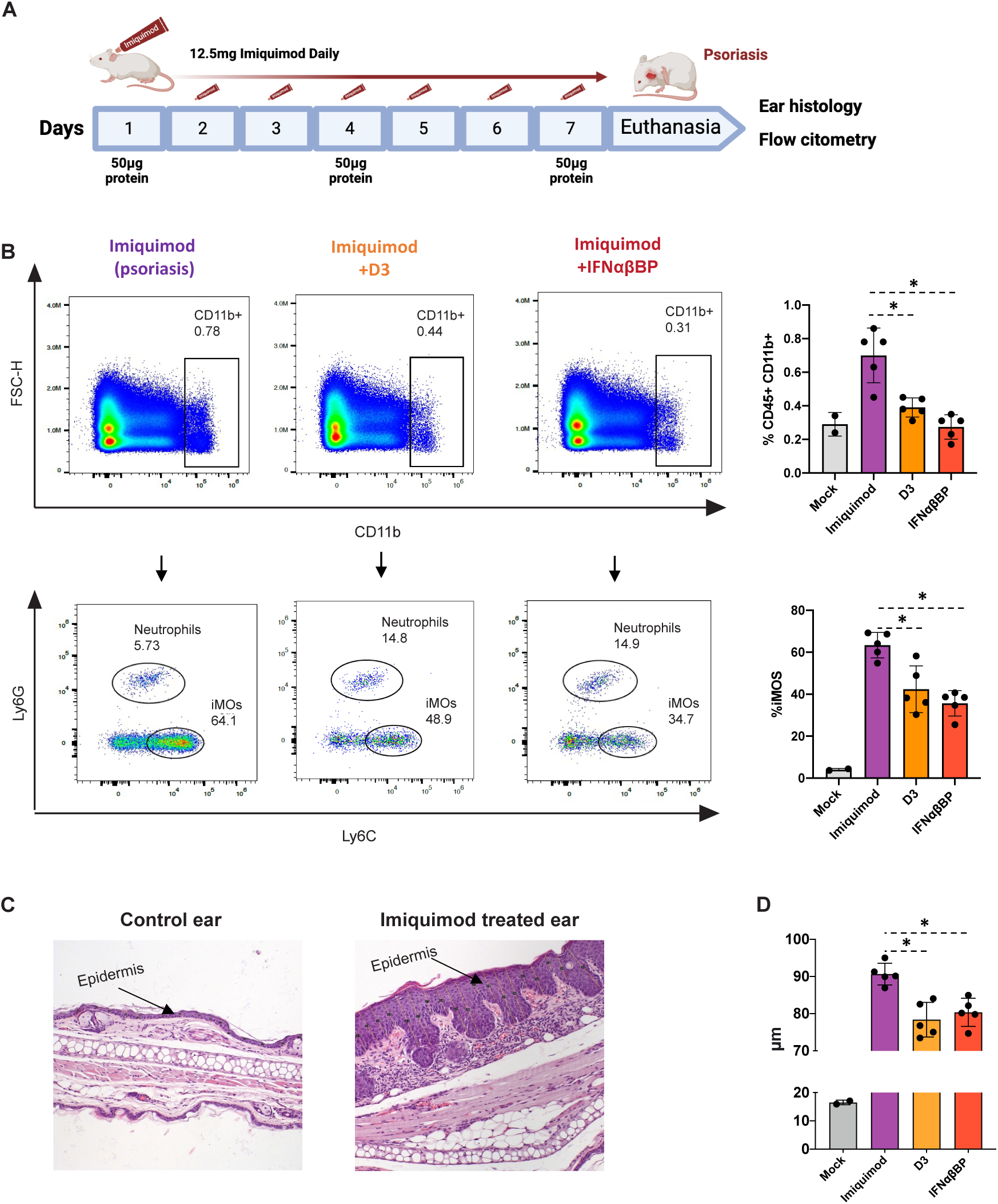
Viral IFN-I inhibitor treatment in a mouse model of IIP. **(A)** 12.5 mg of Aldara cream was applied to each ear of 8-week-old female BALB/c mice during 6 consecutive days. During this time, mice were treated with 50 µg of IFNαβBP, D3 or PBS every 72h. **(B)** Levels of myeloid cells (CD45+ CD11b+) and inflammatory monocytes (CD45+ CD11b+ Ly6C+) in the LNs of IIP mice. **(C)** Examples of healthy and psoriatic ears stained with hematoxylin-eosin. **(D)** Medium length of the epidermal thickening measures grouped by ear (each dot corresponds to the average of 360 measures in a single ear). Results were analyzed with Mann Whitney test. * = p<0.05. N=5 mice for all IIP groups and 2 mice for mock LNs. A representative experiment of two experiments performed is shown.

Treatment with viral IFN-I inhibitors showed clear clinical improvements in the LNs, which were macroscopically enlarged in untreated IIP mice and appeared reduced in treated mice. When analyzed by flow cytometry, a significant reduction in the levels of myeloid cells (CD45+, Cd11b+) and inflammatory monocytes (CD45+, Cd11b+, Ly6C+) was observed in mice treated with the IFNαβBP or D3 (**Fig. 6 B**).

Next, the impact of the treatments in the epidermal thickening of the psoriatic ears was analyzed. IIP is characterized by the marked thickening of the epidermal layer (**Fig. 6 C**), and most studies using this model base their success on reducing this parameter (58,59). Given the great heterogeneity of the epidermal layer, a large number of measures were performed to cover the whole ear surface and estimate the epidermal thickening accurately. Mouse ears treated with the viral IFN-I inhibitors showed a significant reduction of around 10% compared to untreated psoriatic ears (**Fig. 6 D**). In summary, treatment with the viral IFN-I inhibitors reduced the inflammatory state of the LNs and the epidermal thickening in IIP mice.

## Discussion

The pathologies associated with IFN-I dysregulation are still a subject of intense investigation. The IFN-I system stands out for its complexity, with humans encoding 16 different IFN-Is that seem to induce very similar ISGF3 mediated signaling (60). Therefore, it is not surprising that monoclonal antibodies against specific IFN-I ligands have not fully succeeded in treating the multi-factorial, not completely understood interferonopathies.

The IFNαβBP is a unique anti-IFN-I mechanism with broad specificity and high conservation among orthopoxviruses (32,33). In this study, we focused on its unique structural features and showed how D3 directly interacts with the key region of IFN-I that is recognized by the cellular IFNAR2 (45–47). This finding also explains why the IFNαβBP struggles to block mIFNβ (33), as this ligand is an exception that uses IFNAR1 as the high affinity receptor and has been reported to signal independently from IFNAR2 (61,62). Finding this key interaction means that, although the whole IFNαβBP structure seems designed to bind around IFN-I and mask most of its surface, the critical blockade happens through a defined domain that only represents a third part of the full protein. This has remarkable implications for the potential development of IFNαβBP-based therapies, as molecule size is critical for the pharmacokinetics, biodistribution and immunogenicity of the potential therapeutics.

In this line, recombinant IFNαβBP based proteins containing D3 were able to bind diverse IFN-Is and block their antiviral effect, while D1+D2 was unable to do so. Interestingly, D3 alone showed the capacity to inhibit various IFN-Is and was the best hIFNα inhibitor among the designed proteins, making it a molecule of great therapeutic interest. However, the affinity and blocking capacities of the D3-containing proteins was 1-2 orders of magnitude lower than that of the native IFNNαβBP. This suggests that although the key binding happens through D3, all three domains of the IFNNαβBP cooperate to bind and block IFN-I with high affinity.

In addition, defining the critical residues implicated in the blockade of IFN-I may have interesting applications for poxvirus based-vaccines. Some of the safer VACV vaccines against smallpox encoded truncated IFNαβBPs with deletions in D3, like VACV Wyeth, ACAM2000 and MVA (34,37,63). However, it appears that the presence of an inactive IFNαβBP can be beneficial for the induction of immunity against poxviruses, as the elimination of the truncated IFNαβBP in VACV-Wyeth decreases its immunogenicity (64). This is further supported by the IFNαβBP being a natural target for the antibody response, and by the fact that treatment with monoclonal antibodies against the IFNαβBP can cure mice from otherwise lethal mousepox (65), due to the neutralization of the IFNαβBP activity that is required for full ECTV virulence. Therefore, by specifically mutating the key residues that mediate IFN-I blockade, a full but inactive IFNαβBP could be engineered, so that retained its full key immunogenicity while maintaining a good safety profile for the vaccine.

Next, the possibility of transferring the cell surface binding ability to the minimal IFN-I inhibitor D3 was studied. Since the addition of the GAGBD only conferred a small level of binding to the cell surface, it appears that although this specific motif is essential for the cell surface binding ability (35), it is insufficient to fully transfer this ability to D3, probably due to an imperfect folding or insufficient exposure. In addition, only D1+D3 clearly showed the ability to bind GAGs and block IFN-I at the same time.

The potential of D3 to exert anti-IFN-I activity *in vivo* was first proven in the recombinant ECTV infection model, as ECTV D3 and ECTV D1+D3 showed increased virulence in mice compared to ECTV ΔIFNαβBP. However, both recombinant viruses displayed similar virulence and were attenuated when compared to ECTV WT. Although the ECTV infection model has been key to study poxviral immune evasion mechanisms in the past, the dramatic attenuation caused by modifications of the IFNαβBP made it difficult to analyze the immunomodulatory capacity of D3 in this setting. Therefore, we decided to directly study D3 and the IFNαβBP in mouse models of autoinflammatory disease.

The results in PIL mice showed how both IFN-I inhibitors were able to reduce the activation levels of DCs, one of the key therapeutic targets in SLE and a marker for clinical improvement (66,67). Anifrolumab, a specific antibody targeting IFNAR, has been recently approved in the European Union for the treatment of moderate to severe SLE (68,69). The success of this antibody probably relies on its ability to impair signaling of all IFN-I ligands by targeting their common receptor. In this line, a soluble receptor with broad specificity for IFN-I ligands like the IFNαβBP (31,32) represents a promising alternative and a useful tool for combination therapies aimed at the simultaneous blockade of different IFN-I subtypes.

The results in IIP mice showed a marked decrease in the inflammatory state of the LNs and a reduction in the psoriatic ear thickening that appears in this model. Although this reduction was relatively small (around 10%), it has been reported that part of the pathology in IIP mice is independent of the TLR7-imiquimod interaction (70) and thus untreatable with an IFN-I inhibitors. IFN-I upregulation is not the only pathological hallmark in interferonopathies, as some studies suggest that IFN could not be central to the pathology in all cases (71–73). Therefore, the efficacy of an anti-IFN-I therapy will always depend on how much the pathology is in fact due to IFN-I upregulation and how much is related to other functions of the mutated genes. The fact that the immunomodulatory effect was observed on both the skin and LNs of IIP mice despite the IFN-I independent effects of Aldara in this model highlights the therapeutic potential of these viral proteins.

An additional interesting model to explore would be the Trex1 (-/-) mouse model of AGS (74). Trex1 deficiency causes accumulation of ssDNA derived from endogenous retroelements (75), and Trex1 (-/-) mice exhibit dramatically reduced survival due to lethal IFN-driven autoimmune disease (76). As Trex1 deficiency causes AGS syndrome in humans, this mouse model could be used to study the therapeutic potential of viral IFN-I inhibitors in the prototypic type-I interferonopathy.

Notably, during the only previous report that tested the therapeutic potential of the IFNαβBP *in vivo*, the calculated half-life of the injected protein was only 3 hours (43). Therefore, it is possible that therapeutic effects would benefit greatly from increasing the body-residence time of the protein. Chemical modifications like PEGylation can increase the half-life of drugs by several mechanisms (77). In fact, although PEGylation may lead to loss of biological activity, the prolonged body-residence time can usually compensate this effect and result in a better performance *in vivo*, like the case of the PEGylated α-interferon (78). Exploring how different modifications can improve the *in vivo* stability and reduce the immunogenicity of the IFNαβBP and derived molecules will be relevant for the future development of viral immunomodulatory proteins as therapeutics (38,39).

In summary, we have analyzed the interaction between IFNαβBP and IFN-I, identified and modified the minimal IFN-I inhibitor D3 and described its immunomodulatory potential in three different murine models, paving the way for the development of new anti-IFN-I treatments for type-I interferonopathies.

## Materials and methods

### Cells and viruses

L929 (ATCC CCL-1), HeLa (ATCC CCL-2) and BSC-1 (ATCC CCL-26) cells were grown at 37 °C in Dulbecco’s modified Eagle medium (DMEM, Gibco) supplemented with 5% FBS, 2 mM L-glutamine and antibiotics. CHO-K1 (ATCC CCL-61) and the mutants CHO-677 (ATCC CRL-2244) and CHO-745 (ATCC CRL-2242) were maintained in in DMEM-Ham′s F12 (1:1) with similar supplements. High Five insect cells maintained in TC100 medium (Gibco) supplemented with 10% fetal bovine serum for adherent culture or (FBS, Sigma) or Express Five medium (Gibco) for suspension culture.

The ECTV Naval strain (NCBI:txid1651168) was previously described (79) and cultured in BSC-1 cells. Viral stocks were purified by centrifugation through a 36% sucrose cushion, stored in 10mM Tris-HCl pH 9.0 at −80 °C and titrated twice by plaque assay in BSC-1 cell monolayers prior to animal infections. For this, serial dilutions of viral stocks were plated in duplicate on BSC-1 cell monolayers, incubated for 90 min at 37 °C and then replaced with semi-solid CMC DMEM medium supplemented with 2% FBS. Cells were fixed in 10% formaldehyde at 6 days post-infection and plaques were stained with 0.1% (w/v) crystal violet. The Vesicular stomatitis virus strain Cocal (NCBI:txid11276) expressing GFP was kindly provided from Rafael Sanjuan (80), grown in HeLa cells and titrations by plaque assay were performed in BSC-1 cells.

To obtain the ECTV-infected cell supernatants, 10^6^ BSC-1 cells were infected with recombinant ECTVs at multiplicity of infection (MOI) 0.01 pfu/cell. At 72 hpi, cell culture supernatants were collected and inactivated with psoralen a UV light (81). Then, inactivated supernatants were concentrated 80x in Amicon columns (Merck) and stored at -80 °C in single use aliquots

### Expression and purification of proteins

The recombinant baculoviruses expressing the VACV IFNαβBP have been described previously (32,35). The coding sequences for D3, GAGBD+D3, D1+D3 and D1+D2 were amplified by PCR (primers in **Supplementary table S1**) from the sequence of the VACV WR B18R gene (GenBank: D01019.1) and cloned into pAL7, a modified pFastBac1 vector bearing the honeybee melittin signal peptide at the 5′ region and a C-terminal V5-6xHis tag. The generated plasmids pJA7, pJA8, pJA9 and pJA13 were sequenced to confirm the desired modifications and the absence of inadvertent mutations. Then, the plasmids were used to obtain recombinant baculoviruses using the Bac-to-Bac system (Invitrogen). After infection with the recombinant baculoviruses, supernatants from Hi5 cells were concentrated by ultracentrifugation and then buffer exchanged against 50mM phosphate, 300mM NaCl and 10mM imidazole, pH 7.4, prior to affinity-purification of recombinant proteins with Ni-NTA agarose (Qiagen), as described previously (35). Protein purity and quantity were analyzed by Coomasie blue-stained SDS-PAGE and quantified by gel densitometry.

### SPR assays

To determine the affinity constants for the interactions D3, GAGBD+D3, D1+D3 and IFNαβBP were immobilized by amine coupling on CM5 chips (Biacore, GE Healthcare) in order to obtain approximately 350 response units. SPR experiments were performed using a BIAcore X100 biosensor (GE Healthcare). Kinetic analysis for mIFNα was performed in multi-cycle kinetics, injecting each analyte or blank in a separate cycle. For the human IFNs, kinetic analysis was performed in a kinetic titration (single cycle) manner, injecting several times from low to high concentrations with short dissociation times in between, and all injections were analyzed in one sensorgram after a long dissociation time (82). All kinetics were performed at least twice except those for hIFNα, due to the large of amount of cytokine required for these experiments. However, the D3-hIFNα experiment was performed twice to confirm this particularly high affinity.

### Neutralization of IFN-I antiviral activity

The biological assay to measure IFN-I blockade has been described before (32). Here we used the viral-driven fluorescence of a GFP-tagged VSV to estimate infection instead of measuring cell survival. Shortly, L929 (3000 cells/well) or HeLa (5000 cellls/well) cells were seeded in multiwell 96 plates (M96) in DMEM supplemented with 5% FBS the day before the assay. The next day, appropriate dilutions of mIFN⍺ (PBL 12100-1), hIFNβ (R&D 8499-IF/CF) or hIFN⍺ (PBL 11100-1) in DMEM supplemented with 2% FBS and containing increasing amounts of recombinant proteins or infected cell supernatants were prepared and incubated for 1 h at 37 °C before adding them to cells in triplicate. Cells were incubated with the mixture for 16h, allowing free IFN-I to trigger the activation of the antiviral state. For the modified washing assay, cells were incubated with the proteins for 1h before being thoroughly washed and then incubated with IFN-I, so only the proteins that remained attached to the cell surface would be able to block the antiviral effect of IFN-I. Next, cells were infected with GFP-tagged VSV at MOI 2 pfu/cell and viral-driven GFP was measured at 16, 24 and 48 hpi using a FLUOstar Omega (BMG Labtech) plate reader.

### Cell surface binding analysis

To visualize cell surface binding capacity by immunofluorescence, CHO-K1 cells were seeded onto glass coverslips in M24 plates and incubated with 250 nM of purified recombinant viral proteins for 30 min at 4 °C. Cells were then extensively washed with ice-cold PBS and fixed with 4% paraformaldehyde in PBS for 12 min at RT. The membrane permeabilization step was omitted to avoid protein internalization, thus after aldehyde quenching with 50 mM NH_4_Cl for 5 min, cells were incubated with a monoclonal anti-V5 antibody diluted 1:500 (Invitrogen) followed by anti-mouse IgG-A488 (Molecular Probes) diluted 1:1000. Cell nuclei were stained with DAPI (Calbiochem). Images were acquired in a Leica DMI6000B automated inverted microscope equipped with a Hamamatsu Orca R2 digital camera and analyzed with ImageJ.

To quantify cell surface binding capacity by flow cytometry, CHO-K1, CHO-677 or CHO-745 cells were detached with 4 mM EDTA at 37 °C and harvested in PBS. Cells (3×10^5^ per experimental point) were incubated for 30 min with 250 nM of the indicated viral recombinant proteins on ice. Then, cells were extensively washed with FACS buffer (PBS, 0.01% sodium azide and 0.5% bovine serum albumin) and incubated for 30 min at 4 °C with monoclonal anti-V5 antibodies (Invitrogen) diluted 1:500 in PBS, followed by anti-mouse IgG-A488 (Molecular Probes) diluted 1:500 in PBS. Finally, cells were resuspended in 0.3 ml flow cytometry buffer (FC buffer, PBS 1x supplemented with 2% FBS) and acquired in a FACS Canto II flow cytometer (BD Sciences). Data was analyzed with FlowJo v10.9.0 software.

### Construction of recombinant ECTVs

Recombinant ECTVs were generated using ECTV Naval as a parental virus and the previously described transient-dominant selection procedure based on puromycin resistance (83). Briefly, the 5′ (including the signal peptide) and 3′ (including the stop codon) flanking regions of the EVN194 gene (ECTV Naval genome positions 184,714 to 185,726 and 186,710 to 187,457) were inserted into the plasmid pMS30 (83) to produce the plasmid pJA14, that allows recombination into the EVN194 locus. Then, the sequences for D3 and D1+D3 were amplified from pJA7 and pJA9 and subcloned into pJA14 by InFusion Cloning (Takara) to produce pJA15 and pJA17. Then, BSC-1 cells were infected with ECTV-Naval (0.01 pfu/cell) and at 1 hpi transfected with pJA15 and pJA17 using Fugene HD (Invitrogen). When cytopathic effect was complete, cells were harvested and used as inoculum in five consecutive rounds of infection in the presence of 15 μg/ml puromycin (Sigma) monitoring for EGFP expression. ECTV D3 and ECTV D1+D3 were finally isolated by three successive plaque purification steps of white plaques in the absence of puromycin and checked for the desired mutation by PCR (forward primer JA_E194check_1, ttattgctattccatagctacgc; reverse primer JA_E194check_2, ttcagtaataattcagtaatgtatataaaaatgca).

After DNA isolation by treatment with a nuclease mix (DNAsa I, Micrococcal Nuclease S7 and RNAsa A; Roche) and proteinase K (Invitrogen) followed by phenol-chloroform extraction, the presence of viral DNA was confirmed by PCR with the specific oligonucleotides JA_E194check_1 and JA_E194check_2. The contamination with bacterial DNA was discarded by PCR with oligonucleotides targeting the 16S ribosomal RNA gene; 27F: 5’-agagtttgatcmtggctcag-3’, 1492R: 3’-tacggytaccttgttacgact5’. Next, libraries for Illumina sequencing were prepared using Illumina DNA Prep Tagmentation kit following manufactureŕs instructions and NGS sequencing was performed in a MiniSeq system (Illumina) using a 2 x 150 run with the MiniSeq Mid Output kit (300-cycle) (Illumina). The obtained reads were mapped against the corresponding expected reference genome with Bowtie2 aligner software using default parameters and visualized using Integrative Genomics Viewer (IGV). The coverage for both viral genomes was above 500x and the corresponding sequence analysis discarded the presence of additional or non-desirable mutations. Files containing raw reads were deposited at ENA (European Nucleotide Archive) under project number PRJEB94895.

### Virus growth curves

Pre-confluent BSC-1 cells were infected for 1 h at 37 °C at high MOI (5 pfu/cell) or low MOI (0.01 pfu/cell) to analyze single cycle infection and infection progression, respectively. Cells were then washed and fresh medium was added. In the multi-step growth curve, at indicated times post-infection the medium was harvested in DMEM 2% FBS and centrifuged at 1800g for 5 min to pellet detached cells. These cells were combined with infected cells that had been scraped from the plate into 0.5 ml of fresh medium. In the one-step growth curve, cells and media were harvested separately in DMEM 2% FBS at the indicated times. In both cases, samples were frozen/thawed three times and titrated in duplicates on BSC-1 cells as described above.

### Mice

BALB/c ByJ 6–9 weeks old female mice were purchased from Charles River. Mice had access to diet *ad libitum* and were housed in specific pathogen free (SPF) conditions under a 12 h light/dark cycle, and mice were habituated to housing conditions for at least 7 days before the start of experiments.

### ECTV infection of mice

Mice infections were performed with sucrose cushion-purified virus diluted in PBS containing 0.1% BSA. Mice were anesthetized with isoflurane and then infected with ECTV by subcutaneous inoculation into the footpad with 10 μl of virus inoculum containing the indicated infectious doses. Inoculums were back tittered on the day of infection to confirm the inoculated dose. Mice were housed in groups of 5 individuals in ventilated racks under BSL-3 containment facilities equipped with HEPA filters at the CBM Severo Ochoa animal facility. Starting from day 3 post infection, mice were monitored daily for survival, weight loss, footpad inflammation and signs of disease for at least 16 days. To avoid unnecessary suffering, mice bearing a weight loss >20% and disease score >2.5 were euthanized and considered dead by the infection. All animal procedures were approved by the appropriate ethics committees and the legal authorities (PROEX: 241.1).

### Autoinflammatory disease models

For the PIL model, 7-week-old female BALB/c mice were intraperitoneally injected with 0.5ml sterile pristane (Sigma) to induce a lupus-like syndrome. Mice were then intraperitoneally injected with 50 μg of recombinant proteins or PBS on days 2, 5, 8 and 10 after pristane injection. At the experimental endpoint, mice were euthanized and the immune infiltrate in the peritoneal cavity was analyzed by flow cytometry.

For the IIP model, 7-week-old female BALB/c mice were treated daily for 6 consecutive days with 25 mg of cream 5% Imiquimod (Aldara; Meda Pharma) divided among both ears. During this process, mice were also intraperitoneally injected with 50 μg of recombinant proteins or PBS on days 1, 4 and 7. At the experimental endpoint, flow cytometry was performed on the draining LNs, and one ear was fixed for histology.

### Analysis of immune cell populations by flow cytometry

Tissues were specifically preprocessed to isolate cells before flow cytometry. For the PIL model, peritoneal washes were performed by flushing 10 ml PBS in the peritoneal cavity, carefully massaging the abdomens and collecting a minimum of 8 ml of peritoneal liquid containing cells in suspension. For the psoriasis model, axillar and subiliac LNs were aseptically harvested, pooled and mechanically disaggregated by filtering through 100 μm nylon cell strainers (Falcon).

For the staining protocol, cells were transferred to a M96 V-bottom plate and washed twice with PBS. Then, cells were stained for viability by incubating for 20 min at 4 °C with Ghost Dye-v540 (ThermoFisher) in PBS. After washing twice with flow cytometry buffer (FC buffer, PBS 1x supplemented with 2% FBS), cells were incubated for 15 min with Fc block (BD 553142).) to reduce non-specific staining. Then, cells were incubated with surface antibodies for 20 min at 4 °C. Finally, cells were washed and resuspended in FC buffer and acquired in an Aurora spectral flow cytometer (Cytek). The acquired data was analyzed in Flowjo v10.9.0 software.

### Antibodies

From BD: BV421 anti-Ly6C (BD-562727), RB545 anti-B220 (BD-756542), PE anti-CD11b (BD-557397), PE-CF594 anti-CD86 (BD-567592), APC anti-CD11c (BD-550261), BB700 anti-CD49b (BD-568015), R718 anti-CD80 (BD-752045), BV50 anti-CD3 (BD-746988), PE-Cy7 anti-CD27 (BD-563604), APC-Cy7 anti-CD4 (BD-565650), RB780 anti-Ly6G (BD-569153). From Biolegend: BV605 anti-MHC-II (Biolegend-107639), APC/Fire810 anti-CD45 (Biolegend-103173).

### Histology

After euthanizing the IIP mice, whole ears were aseptically removed and fixed in formalin for 2h at RT and then overnight at 4 °C. The histological cuts and hematoxylin-eosin staining were prepared by the histology service at Centro Nacional de Biotecnología (Madrid), sampling three different sections of the ears separated by approximately 700 μm. To estimate the epidermal thickening, 8 different pictures were taken for each of the three ear sections, and 15 individual measures of the epidermis were made in each picture. Images were captured using an Olympus microscope BX41, 10x objective, with an Olympus camera DP-70 (Olympus Denmark A/S). Epidermal thickness was quantified using ImageJ.

### In silico structure analysis

Protein structures were modeled using Alphafold v2 (84) in a publicly available server (85). The heparin-protein docking was performed using the server ClusPro (86). The resulting structures were visualized and analyzed in Chimera X (87), overlapping over the crystal structure PDB:3OQ3 to visualize the potential binding of IFN-I.

### Statistical analysis

Data was analyzed using GraphPad Prism (version 8) software. Footpad swelling and % of initial weight data analyzed using multiple t-tests with false discovery rate (FDR) Q = 1%. For the data related to IFN neutralization assays and flow cytometry, medians were compared with the non-parametric Mann–Whitney U-test at a significance level α=0.05. For the ear thickening data, Mann–Whitney U-test was performed after grouping the measures by mouse.

## Acknowledgements

This research was funded by the Spanish Ministry of Science, Innovation and University grants PDC2022-133701 and RTI2021-128580OB-I00. F.J.A.M. and I.A.S. were funded by PhD Studentships FPU 19/04346 and FPI PRE2019-090941, respectively.

## Author Contributions

Conceptualization and supervision, A.A. and B.H.; investigation, F.J.A.M, I.A.S, P.D. and A.S.P.; writing—original draft preparation, F.J.A.M.; funding acquisition, A.A. All authors have read and agreed to the published version of the manuscript.

## Notes

### Competing Interest Statement

The authors have declared no competing interest.

