## Supplementary Material for "Viral soluble decoy receptor as a novel treatment for IFN-I triggered diseases"

**Table S1. Oligonucleotides.** List of the oligonucleotides used, indicating the cloning strategy they were used for (restriction enzyme target or In Fusion), their sequence and the target plasmid they were used to construct. In the sequences, specific restriction enzyme sites appear underlined.

| **Oligonucleotide** | **Strategy** | **Sequence (5'-> 3')** | **Plasmid generated** |
|---|---|---|---|
| JA_D3F_3 | BamHI | gcgggatcccacaggtttaaactaatactagatcc | pJA7 |
| JA_D3_2 | NotI | gcggccgctctccaatactactgtagttgtaaggg | pJA7 |
| JA_Sticker_F1 | BamHI | gcgccatggcaggatccaaatggg | pJA8 |
| JA_Sticker_R1 | SalI | gcggaattcgtcgacggaaccacc | pJA8 |
| JA_D3F_2 | SalI | gcggtcgaccacaggtttaaactaatactagatcc | pJA8/9 |
| JA_D1_F | BamHI | gcgggatccatagacatcgaaaatg | pJA9/13 |
| JA_D1_R | SalI | cgcgtcgacttttctaatatgagatctaactataccc | pJA9 |
| JA_D2_R2 | NotI | cgcgcggccgctcctgtggtcttgtgacgg | pJA13 |
| JA_E194FL_R | BamHI | gcgggatccggcgtagctatggaatagcaat | pJA14 |
| E194del1 | SacI | gccgagctcgatgagcgtcgcatgatcgt | pJA14 |
| JA_E194FR_F_3 | BamHI | gccggatcctaaatacacaatgcatttttatatacattact | pJA14 |
| E194del4 | PstI | gccctgcagtccgcgtcatctacatctgat | pJA14 |
| JA14F_2 | In Fusion | taaatacacaatgcattttt | pJA15/17 |
| JA14R_2 | In Fusion | ggcgtagctatggaatagca | pJA1517 |
| 15F_2 | In Fusion | ttccatagctacgcccacaggtttaaactaatactagatc | pJA1517 |
| 15R_2 | In Fusion | tgcattgtgtatttactccaatactactgtagttgtaagg | pJA15 |
| 17F_2 | In Fusion | ttccatagctacgccatagacatcgaaaatgaaatcacag | pJA17 |

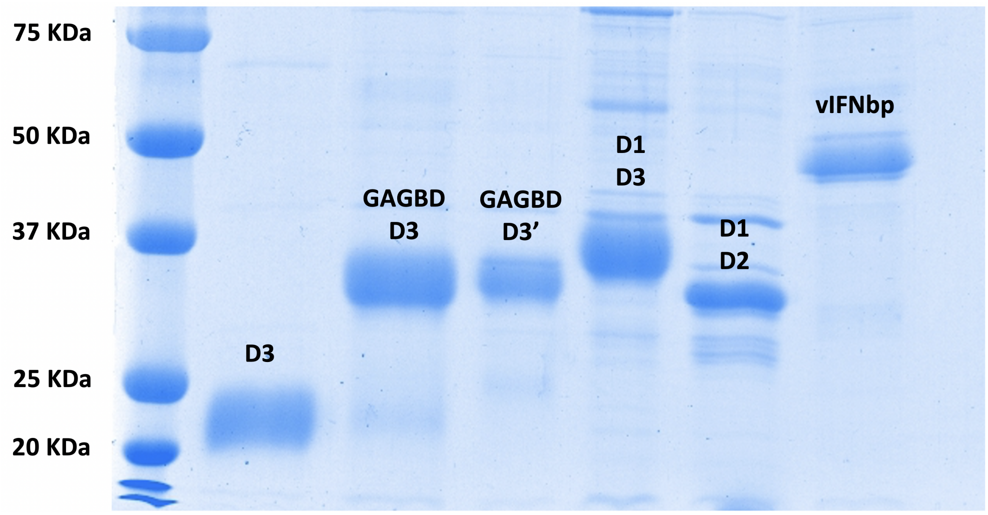


**Figure S1**. SDS-PAGE and Coomasie Blue staining of 1 μg of each IFNαβBP-based recombinant protein expressed in the baculovirus system.

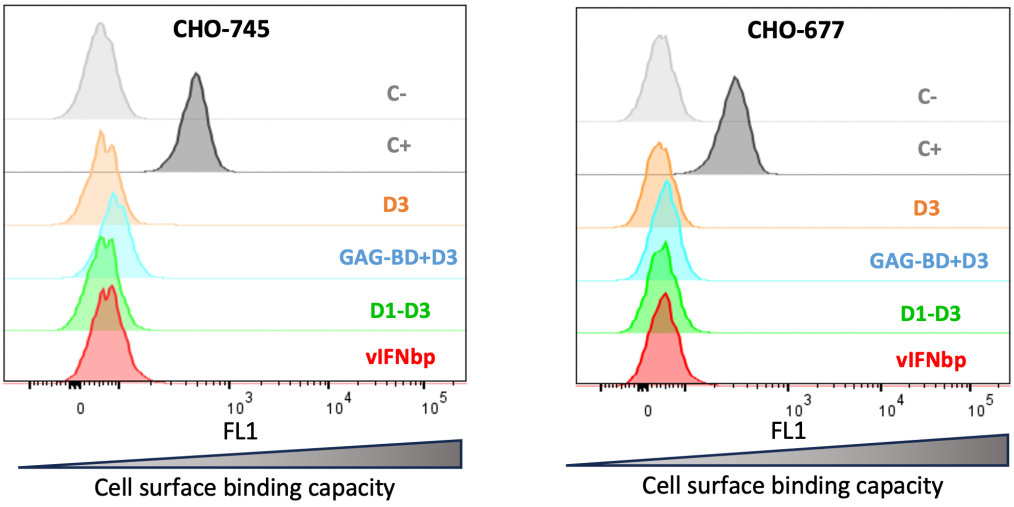


**Figure S2**. Cell surface binding of recombinant proteins to CHO-745 (lacking the GAGs) and CHO-677 (with a specific depletion on heparin) cells, analyzed by flow cytometry. Mpox virus A41 protein is shown as a positive control. A representative experiment from a minimum of two experiments performed is shown

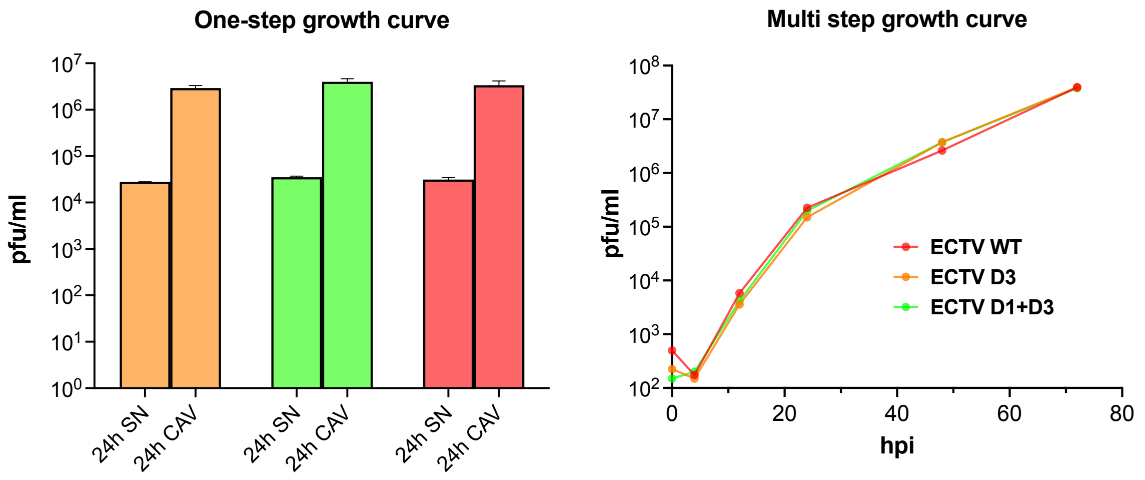


**Figure S3**. **Replication of recombinant ECTVs in BSC-1 cells.** Viral yield at 24 hpi with high MOI (10 pfu/cell) or at several time points during 72h after infection with low MOI (0.01 pfu/cell). SN: virus in supernatant, CAV: cell-associated virus.
